# Characterizing bilateral M1 excitability changes after learning a novel spatiotemporal coordination pattern

**DOI:** 10.64898/2026.09.25.754390

**Authors:** Yiyu Wang, Deanna M. Kennedy, Madison M. Weinrich, Yuming Lei, David L. Wright, John J. Buchanan

**Affiliations:** Department of Psychology, Princeton University, Princeton, New Jersey, United States; Department of Kinesiology and Sport Management, Texas A&M University, Texas, United States; Department of Neurosurgery, Center for Neural Systems Restoration, Houston Methodist Research Institute, Houston, Texas, United States

**Keywords:** Bimanual coordination, Unimanual coordination, Primary Motor Cortex, Interhemispheric inhibition

## Abstract

Bimanual coordination is essential for many skilled everyday actions and requires maintaining specific spatiotemporal relationships between the two hands according to task demands. However, the contribution of the primary motor cortices (M1s) to learning such relationships is only partially understood. Conventional bimanual tasks necessarily engage both M1s in controlling the moving hands, making learning-related changes difficult to distinguish from those associated with movement execution. To reduce this effector-related confound, we used a unimanual and bimanual rhythmic coordination paradigm in which participants learned the same novel 90° relative-phase relationship by coordinating either hand with a virtual partner and compared these conditions with conventional bimanual practice. In three experiments, performance accuracy and stability improved during practice and improved further after a 6-h retention interval with comparable time-dependent changes across unimanual and bimanual trained groups. In Experiment 2, motor-evoked potentials measured before and after training revealed an overall increase in corticospinal excitability across both M1s under unimanual and bimanual training. In Experiment 3, bidirectional dual-site paired-pulse stimulation revealed effector-dependent changes in interhemispheric inhibition, in which bimanual practice produced no reliable directional modulation, whereas unimanual practice produced asymmetric changes that depended on the trained hand. Neither corticospinal excitability nor interhemispheric inhibition showed a consistent relationship with behavioral improvement. Together, these findings demonstrate comparable behavioral learning of a novel spatiotemporal coordination relationship across bimanual and unimanual contexts, while showing that behavioral improvement is not straightforwardly reflected in M1 neurophysiological changes. Causal intervention studies are therefore needed to determine whether and when M1 contributes directly to acquisition of novel coordination relationships.

## Introduction

Many everyday activities depend on bimanual actions that require continuous coordination between the two hands to maintain specific spatiotemporal relationships. Examples range from driving a manual-transmission vehicle and tying shoelaces to playing the piano. The ability to coordinate the hands with precision is important for completing many daily activities that support functional independence to maintain quality of life. Understanding the behavioral and neural mechanisms underlying bimanual coordination is consequently important for explaining how the nervous system organizes functionally relevant everyday actions.

Coordination dynamics theory describes the emergence of spatiotemporal patterns of coordination through self-organizing processes (Kelso, 1995). The Haken–Kelso–Bunz model provides a mathematical account of the dynamics of rhythmic bimanual actions, using relative phase (Φ) as an order parameter that captures the spatiotemporal relationship between movements of the two hands (Haken et al., 1985). A 90° relative phase pattern occurs when one hand leads the other by one-quarter of a movement cycle. Unlike the intrinsically stable in-phase (Φ = 0°) and anti-phase (Φ = 180°) coordination patterns, the 90° pattern is not an intrinsic attractor and is inherently unstable, tending to drift toward one of the more stable coordination states (Haken et al., 1985). Studies have demonstrated that the novel 90° relative phase pattern can be acquired and stabilized through practice (Kovacs et al., 2009b, 2009c; Zanone & Kelso, 1992), making it a useful coordination task for investigating how the dynamics of a bimanual motor skills are acquired and retained.

The primary motor cortex (M1) has been widely implicated in motor learning and motor memory consolidation that stabilizes newly acquired motor memories through the resting period after the completion of the initial practice (Krakauer & Shadmehr, 2006; Shadmehr & Brashers-Krug, 1997). This is supported by evidence showing that neural activities at M1 are linked to behavioral changes during motor skill acquisition and subsequent motor memory consolidation (Bhattacharjee et al., 2021). For example, disrupting M1 activity immediately after practice can destabilize newly acquired motor memories (Muellbacher et al., 2002), and post-training modulation of M1 activity has been shown to influence the consolidation of motor sequence learning (Robertson et al., 2005; Tunovic et al., 2014; Wang, Huynh, et al., 2024). Evidence from sensorimotor adaptation further implicates M1 in motor-memory formation, although its specific contribution appears to depend on the task and the stage of learning(Herzfeld et al., 2014; Hunter et al., 2009; Richardson et al., 2006).

In contrast, evidence regarding the contribution of M1 to the acquisition and consolidation of novel bimanual coordination patterns, as indexed by changes in corticospinal excitability, has been less consistent. Nomura et al. (2016) showed that performing a 90° bimanual relative phase pattern was accompanied by increased corticospinal excitability in the dominant M1 of right-handed participants, with a greater increase than observed during in-phase and anti-phase coordination (Nomura et al., 2016). This finding suggests that corticospinal excitability is sensitive to the greater control demands associated with producing a more complex coordination pattern. Consistent with this observation, Park et al. (2020) reported increased dominant-M1 corticospinal excitability following training of the same 90° pattern (I. Park et al., 2020). However, the magnitude of this increase was not associated with consolidation-related improvements in relative-phase accuracy. Moreover, increases in corticospinal excitability are not unique to novel coordination patterns, as excitability changes have also been observed during or following performance of more intrinsic or symmetric bimanual actions (Byblow et al., 2012; Nomura et al., 2016). Together, these findings leave the functional significance of M1 excitability changes for bimanual coordination.

Several factors may contribute to these mixed findings. First, most previous studies have assessed corticospinal excitability in only one M1, typically the dominant hemisphere. Because bimanual coordination requires simultaneous control of both hands, both M1s are likely involved in performance, and measurements restricted to one hemisphere may therefore provide an incomplete account of learning-related neurophysiological changes. Second, changes in corticospinal excitability following practice may reflect multiple processes associated with task performance. In conventional bimanual paradigms, acquisition of the spatiotemporal coordination pattern occurs concurrently with repeated execution of movements by both hands, making it difficult to distinguish plasticity associated with learning the coordination relationship from that associated with movement execution itself. This limitation motivates coordination paradigms that can dissociate acquisition of the spatiotemporal pattern from the specific motor commands used to implement it. Third, bimanual coordination may depend not only on excitability within each M1 but also on interactions between the two hemispheres. Coordinating the two hands requires integration of motor commands across bilateral motor networks, supported in part by transcallosal communication between motor cortices (Cunningham et al., 2017; de Oliveira, 2002; Fujiyama et al., 2016; Mayston et al., 1999; Serrien et al., 2003). Interhemispheric inhibition (IHI), measured using dual-site TMS, provides a neurophysiological index of inhibitory interactions between the two M1s and has been shown to vary with motor performance and learning and to relate to interlimb transfer (Paparella et al., 2023; Perez et al., 2007). Given the importance of interhemispheric interactions for bimanual control (Carson, 2005; Gooijers & Swinnen, 2014), changes in IHI may therefore contribute to the acquisition of accurate bimanual coordination. This study sought to identify if IHI changes underpin behavioral performance outcomes that emerge when an unstable rhythmic coordination pattern stabilizes with practice.

Unimanual coordination with a computer-programmed virtual partner provides a useful approach for dissociating neural processes associated with learning from those associated with executing the same spatiotemporal coordination pattern (Kelso et al., 2009). To produce a virtual partner relative phase pattern, the participant must continuously synchronize the movement of one hand with an externally generated visual signal (Kelso et al., 2009; Snapp-Childs et al., 2011). Execution of unilateral hand movements is predominantly supported by the contralateral motor cortex, whereas ipsilateral M1 involvement is weaker, yet can increase with task complexity (Koeneke et al., 2004). Training-related changes observed in M1 ipsilateral to the active hand are less readily attributable to direct motor output and may instead reflect broader bilateral processes associated with acquiring and representing a novel coordination pattern. Moreover, assigning the same coordination relationship separately to the left and right hands provides complementary conditions for determining whether learning-related changes are lateralized to the hemisphere controlling the active hand or shared across both M1s. In this way, the unimanual paradigm provides a means of probing how the two hemispheres contribute to acquisition of a common spatiotemporal coordination pattern while minimizing the confound introduced by simultaneous execution of two voluntary hand movements.

Building on this rationale, the present study examined whether learning the same 90° relative phase pattern under different effector configurations is supported by common or distinct neurophysiological processes within the bilateral motor system. Specifically, we asked two questions: 1) Are practice-induced changes in corticospinal excitability in the left and right M1 related to improvements in coordination accuracy and stability during practice and across a subsequent 6-hour post-training consolidation interval; and 2) Are changes in bidirectional IHI similarly associated with online and offline (post-training) behavioral changes. Experiment 1 established the validity of the unimanual coordination paradigms for addressing these questions by determining whether practice with either hand produced behavioral signatures comparable to those observed during bimanual coordination, including acquisition, and post-training improvement. Establishing these shared behavioral characteristics was critical for ensuring that subsequent neurophysiological comparisons reflected different neural implementations of a common coordination structure rather than learning of fundamentally different task-specific solutions. Experiment 2 used single-pulse TMS to determine whether acquisition of this common coordination structure produced similar or distinct changes in corticospinal excitability across the two M1s. Experiment 3 used dual-site paired-pulse TMS to determine if the same learning conditions differentially reorganized inhibitory interactions between the two hemispheres. Previous intracranial, fMRI, and EEG studies have demonstrated hemispheric asymmetries in motor control, with the dominant left hemisphere in right-handed individuals showing broader bilateral involvement during movement preparation and execution (Merrick et al., 2022; Serrien et al., 2003; Serrien et al., 2006; Swinnen et al., 2010). Based on this evidence, we hypothesized that learning across all three training conditions would preserve or strengthen the functional influence of the dominant left M1, reflected by greater left-hemisphere corticospinal modulation and/or a shift in IHI favoring left M1. In contrast, modulation of right M1 was predicted to depend more strongly on effector configuration, with greater involvement during left-hand and bimanual practice than during right-hand practice.

## 2. Method

### Participants

A total of 112 participants (age = 21.42 ± 2.48 years; 72 female) were recruited from the research pool at Texas A&M University. The Edinburgh Handedness Inventory confirmed that all participants were right-hand dominant (Oldfield, 1971). The Institutional Review Board at Texas A&M University approved the procedures, and all participants provided written informed consent before participation. For experiment 1, thirty-eight participants were randomly assigned to the bimanual (n = 12), unimanual-left (n = 12), or unimanual-right (n = 14) group. In Experiment 2, thirty-eight participants were randomly assigned to the bimanual (n = 13), unimanual-right (n = 13), or unimanual-left (n = 12) group. In Experiment 3, thirty-six were randomly assigned to the bimanual (n = 12), unimanual-left (n = 12), or unimanual-right (n = 12) group. Each experiment consisted of subjects naïve to the task.

### Apparatus

The experiments measured index-finger abduction–adduction movements in the horizontal plane while participants trained with a 90° relative phase pattern. Three training contexts were used: bimanual, unimanual-left, and unimanual-right. Bimanual coordination required participants to synchronize rhythmic actions produced by the left and right index fingers. Unimanual-left and unimanual-right coordination required participants to synchronize rhythmic actions of a single index finger with a computer-controlled virtual partner generated by a sine-wave function oscillating along the *x*- or *y*-axis (Figure 1).

**Figure 1.**
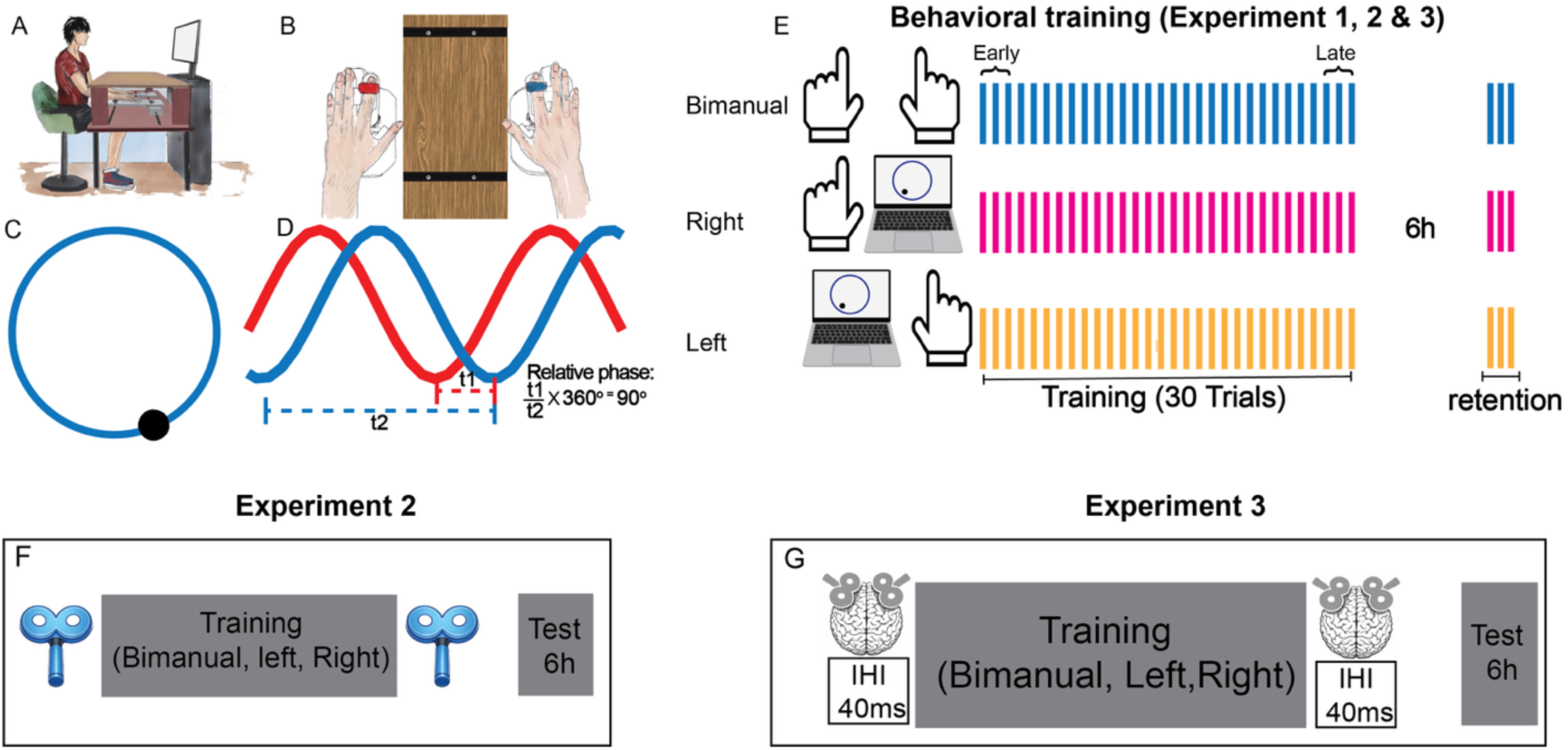
A) Experimental setup. B) Position of hands beneath the countertop when locked into potentiometers measuring finger abduction-adduction movements. C) Lissajous feedback display with a cursor superimposed on circle template representing the 90° relative phase pattern. D) Example of time series illustrating the 90° pattern with one signal a quarter cycle ahead of the other. E) Demonstration of the experimental training design: bimanual (blue), unimanual right (pink), and unimanual left (yellow) training group, with each vertical line representing a 20-second trial. F) The timeline for TMS application in Experiment 2. G) The timeline for IHI stimulation in Experiment 3. Note, behavioral training protocol is used consistently across the three experiments.

Participants sat at a table with seat and tabletop height adjusted individually so that both hands rested comfortably on the tabletop, approximately shoulder-width apart, with the palms facing down (Figure 1A). A custom finger-constraint apparatus was fitted to each hand to allow comfortable abduction–adduction of the index finger while minimizing movement of the other digits (Figure 1B). The index finger was inserted into a slot mechanically coupled to a potentiometer that continuously recorded finger position (Figure 1B). After participants adopted a comfortable posture, their index fingers were secured with straps to prevent sliding within the slots, and the thumb and non-task fingers were stabilized using finger barriers. A wooden countertop occluded the participants’ view of their hands throughout the task (Figure 1A).

Potentiometer signals were recorded with an A/D converter (NI USB-6210, National Instruments, Austin, TX, USA), sampled at 100 Hz, and stored on a PC for offline analysis in MATLAB (R2022a, MathWorks, Natick, MA, USA). Visual feedback was presented on a 27-inch monitor positioned directly in front of the participant. A circle template and a cursor displayed as a Lissajous plot provided online feedback about the target 90° relative phase pattern and about the participant’s own movement in the bimanual and unimanual conditions (Figure 1C). The virtual partner oscillated sinusoidally at a constant frequency of 0.6 Hz. The cursor trajectory displayed on the screen was determined by the coordination between the two hands in the bimanual task, and the coordination between one hand and the virtual-partner in the unimanual tasks (Figure 1D).

### Behavioral task: Experiments 1, 2, and 3

For the bimanual group, the screen cursor was controlled by the two potentiometer signals, with each hand contributing one dimension of cursor motion. The movement of one finger was mapped to the cursor’s *x-axis* and the other to the *y-axis*, such that cursor position (*x, y*) at each time point was defined by the combined signals. To avoid systematic effects of axis assignment, the mapping between hand (left vs. right) and cursor axis (*x* vs. *y*) was pseudorandomly counterbalanced across participants. In the unimanual-left and unimanual-right groups, cursor position at each time point was defined by the signal produced by the active hand together with that of the virtual partner, and this hand–virtual partner mapping was likewise assigned pseudorandomly to the *x*- or y-axis. The virtual partner moved the cursor along its assigned dimension at a frequency of 0.6 Hz per cycle (a cycle is defined as a sine wave from 0 to 2π). The goal for all groups was to produce the 90° relative phase pattern by coordinating the two fingers or a finger with the virtual partner by moving the cursor along the Lissajous display (Figure 1C–D). In the bimanual group, participants performed the coordinated movements at a self-paced frequency and were reminded to maintain a constant speed across trials.

At the start of the training session, each participant completed a 20-s familiarization trial in which they produced in-phase coordination (ϕ = 0°) to learn the mapping between hand movements and the corresponding direction of cursor motion. Participants then completed 30 training trials of the 90° pattern with concurrent cursor/Lissajous template feedback. Each training trial lasted for 20-s and was followed by a 2-s rest interval. The first three training trials were designated the Early phase and the last three the Late phase; the difference between the Early and Late phases indexed online improvement during acquisition. A retention test of three trials was administered 6-h after the end of training to assess recall of the practiced 90° pattern.

### TMS measurement: Experiments 2 and 3

TMS was delivered using a DuoMAG MP-Dual stimulator (Brainbox Ltd., UK) with monophasic pulses. Stimulation was applied through a 70-mm figure-of-eight coil held at approximately 45° to the midline, with the handle pointing laterally and posteriorly to induce a posterior–anterior current in the cortex (Di Lazzaro et al., 2004). Coil position and orientation were monitored using a frameless stereotaxic neuronavigation system (Brainsight, Rogue Research, Canada) to ensure consistent targeting throughout the session.

For each participant, the M1 “hotspot” was identified separately for the left and right first dorsal interosseous (FDI) muscles. The hotspot was the scalp location that reliably elicited MEPs in the target muscle, measured using surface electromyography (EMG). During the hotspot search, stimulation was delivered while EMG responses were monitored, and a candidate site was accepted as the hotspot if it produced MEP amplitudes greater than 0.05 mV in at least 5 of 10 consecutive trials at the lowest achievable stimulus intensity. Once identified, the hotspot coordinates were saved and the neuronavigation system was used to guide subsequent stimulation. Stimulation intensity was then set to the minimum level that elicited MEPs of at least 1 mV in at least 5 of 10 consecutive trials, and this intensity was used for the experimental TMS pulses.

### Corticospinal excitability: Experiment 2

Because coordination between the hands is not equivalent to the linear addition of two independent one-handed actions (Kelso et al., 1979a, 1979b; Ryu & Buchanan, 2004; Sherwood & Nishimura, 1999), we hypothesized that bimanual and unimanual practice would modulate corticospinal excitability differently. Experiment 2 was designed to test this hypothesis by using TMS to assess corticospinal excitability before and after training (Figure 1F). Twenty single TMS pulses were delivered to left and right M1, and corticospinal excitability was quantified as the peak-to-peak amplitude of the MEP recorded from the FDI muscle contralateral to the stimulated hemisphere, with larger amplitudes indicating greater corticospinal excitability.

### Interhemispheric inhibition (IHI): Experiment 3

To examine whether unimanual and bimanual practice differentially modulated interhemispheric interactions, IHI was assessed a with dual-site paired-pulse TMS procedure before and after training (Harris-Love et al., 2007) (Figure 1G). IHI was measured at a 40-ms interstimulus interval (ISI) in both directions between left M1 and right M1. This design tests whether training-related changes in coordination performance were accompanied by direction-specific changes in inhibitory interactions between the hemispheres. The same DuoMAG MP-Dual stimulator and coil orientation described above were used, except that two 30-mm figure-of-eight coils replaced the single 70-mm coil to accommodate simultaneous bilateral stimulation.

Surface EMG was recorded from the FDI muscle, and participants were instructed to keep both hands relaxed throughout testing. IHI was measured bidirectionally by pairing a conditioning stimulus (CS) over one M1 with a test stimulus (TS) over the opposite M1 (CS_right_ + TS_left_; CS_left_ + TS_right_), with the order of directions counterbalanced across participants. Because the test stimulus was always delivered to the hemisphere receiving the inhibition, each direction was identified by tested hemisphere: the inhibition exerted by left M1 on right M1 was assessed when right M1 was tested, and the inhibition exerted by right M1 on left M1 was assessed when left M1 was tested. For each direction, three stimulus conditions were delivered: unconditioned TS-only trials and conditioned CS–TS trials with an interstimulus interval (ISI) of 40 ms. Each condition comprised 10 trials, yielding 30 trials per direction. The sequence of conditions was randomized by computer, with a 7-s interval between trials.

TS intensity was set separately for each hemisphere as the minimum intensity that elicited MEPs of at least 1 mV in at least 5 of 10 consecutive trials in the contralateral FDI muscle. CS intensity was set to 120% of the resting motor threshold for the stimulated hemisphere. IHI was quantified as the ratio of the conditioned to the unconditioned MEP peak-to-peak amplitude for the same TS site and direction, with smaller ratios indicating stronger inhibition.

#### Data reduction and analyses

All data computation and reduction were performed in MATLAB (R2022a). A low-pass Butterworth filter (10-Hz cutoff) was applied to the recorded potentiometer time series representing finger abduction–adduction motion. The filtered time series were detrended to identify zero-crossing points, which were used to partition the series into half-cycle segments, and each segment was normalized to a range from −1 to 1.

### Coordination Accuracy

Continuous relative phase (*φ*) between the two coordinated components (i.e., the left and right index fingers in the bimanual condition, or the active finger and the virtual partner in the unimanual conditions) was computed for each trial. For each signal, the phase angle at every time point was derived from the normalized displacement and velocity using the position–velocity phase plane method: 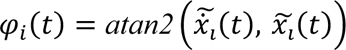, *i* = 1,2, where 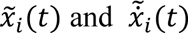 are the within-trial normalized displacement and velocity of component *i*. The continuous relative phase was then calculated as the point-by-point difference between the two-phase angles, *ϕ_rel_(t)* = *ϕ_1_(t) – ϕ_2_(t).* Given that the signal was sampled at 100 Hz for 20 s, each trial produced 2000 instantaneous relative phase values. The initial 2-s (200 samples) of each trial was discarded prior to analysis to remove transient effects associated with movement onset. Coordination accuracy was quantified as the proportion of time during which relative phase remained within a ±30° bandwidth of the target 90° relative phase pattern. This measure, hereafter referred to as BW30, reflects the percentage of trial time during which participants maintained the target pattern within the specified tolerance range, with larger values indicating greater accuracy. BW30 was used in all three experiments.

### Coordination Variability

Coordination variability was quantified using circular statistics applied to the distribution of relative phase values within each trial (Burgess-Limerick et al., 1991; Mardia, 1972). A mean vector length *R* was calculated as a measure of dispersion of the continuous relative phase series (*φ*) with variability computed as *1-R*. This measure captures the dispersion of phase values on the unit circle. Smaller values of *1-R* indicate tighter clustering around the mean phase (greater coordination stability), whereas larger values indicate greater variability, ranged from 0 to 1. The coordination variability measure was used in all three experiments.

### Movement Frequency

Movement frequency was determined using frequency-domain analysis of the displacement signals. For each trial, the displacement time series was mean-centered, and the power spectrum was computed via fast Fourier transform (FFT). The dominant movement frequency (*f*_dom_) was defined as the frequency at which the power spectrum reached its maximum within a physiologically plausible band of 0.5–3 Hz. In the bimanual condition, the task required participants to produce isofrequency (1:1) coordination between the two hands at a 90° relative phase; accordingly, *f*_dom_ was computed separately for each hand and the two values were averaged to yield a single estimate per trial. In the unimanual-left and unimanual-right conditions, *f*_dom_ was associated with the active hand; ideally matching the movement frequency of the virtual partner at 0.6 Hz. Movement frequency was quantified in all three experiments.

### Bilateral corticospinal excitability

For each hemisphere, 20 MEPs were collected at rest before (pre-training) and after (post-training) completion of the behavioral training protocol. MEP amplitudes were averaged within each time point to yield a mean excitability value per hemisphere. Training-induced changes in corticospinal excitability were then computed as the difference between post- and pre-training mean amplitudes: ΔMEP = MEP_post_ – MEP_pre_. Positive ΔMEP values indicate an increase in excitability following training. This approach allowed assessment of bilateral changes in corticospinal output as a function of practice condition. Corticospinal excitability was quantified in Experiment 2 only.

### Interhemispheric Inhibition (IHI)

IHI was quantified using the dual-site paired-pulse TMS protocol by comparing conditioned and unconditioned MEPs. For each stimulation direction (CS_right_ + TS_left_; CS_left_ + TS_right_), mean peak-to-peak MEP amplitudes were computed separately for unconditioned trials, in which the TS was delivered alone, and for conditioned trials, in which the CS was delivered to the contralateral M1 40 ms before the TS. IHI was defined as the ratio of the conditioned to unconditioned MEP amplitude, IHI = (MEP conditioned+test/MEP test), with smaller ratios (closer to 0) indicating stronger interhemispheric inhibition, whereas ratios closer to 1 indicate weaker inhibition (Figure 7A).

To assess training-related changes in interhemispheric inhibition, IHI values were computed before (pre-training) and after (post-training) the behavioral protocol, and the change in IHI was calculated as: ΔIHI = IHI_post_ – IHI_pre_. Because IHI is expressed as a ratio in which smaller values reflect stronger inhibition, positive ΔIHIvalues indicate a reduction in inhibition following training (i.e., disinhibition), whereas negative values indicate an increase in inhibition (Harris-Love et al., 2007). This approach allowed for the assessment of direction-specific changes in interhemispheric communication as a function of training. IHI was quantified in Experiment 3 only.

#### Statistics

All analyses were conducted in MATLAB (R2022a, MathWorks) using the Statistics and Machine Learning Toolbox. Linear mixed-effects models (LMMs) were fit (with the fitlme routine) using restricted maximum likelihood estimation and Satterthwaite’s approximation for degrees of freedom. Effect sizes are reported as partial *eta squared* (*η*^2^). Significance was evaluated at *α = .05* with results between *.05* and *.10* described as marginal. Planned pairwise comparisons of significant main effects or interactions were obtained by re-estimating the model with different reference conditions; reported coefficients reflect the resulting estimated mean differences. One-sample and paired *t*-tests were performed as were correlations using the ttest and corr functions in MATLAB.

For each experiment, coordination accuracy (*BW30*), coordination variability (*1-R*), and movement frequency (*f_dom_*) were each analyzed using a linear mixed-effects model with Group (bimanual, unimanual-left, unimanual-right) as a between-subjects factor and Time (Early, Late, 6-h retention) as a within-subjects factor, with a random intercept for subject. Because both unimanual groups were required to synchronize with a virtual partner moving at 0.6 Hz, we additionally conducted one-sample *t*-tests comparing movement frequency in the unimanual-left and unimanual-right groups against this target value. Movement frequency was analyzed as a potential control variable, given that faster movement speeds can independently affect coordination accuracy and variability.

ΔMEP was analyzed using a linear mixed-effects model with Group (bimanual, unimanual-left, unimanual-right) as a between-subjects factor and Hemisphere (left M1, right M1) as a within-subjects factor, with a random intercept for subject. The model intercept was inspected to assess whether ΔMEPs changed reliably across all groups and hemispheres. To follow up on condition-specific patterns, one-sample *t*-tests were conducted within each Group × Hemisphere condition to determine whether ΔMEP differed from zero.

ΔIHI was analyzed using a linear mixed-effects model with Group (bimanual, unimanual-left, unimanual-right) as a between-subjects factor and Direction (CS_right_ + TS_left_; CS_left_ + TS_right_) as a within-subjects factor, with a random intercept for subject. To decompose significant interactions, we conducted planned pairwise comparisons by re-estimating the model with different reference conditions, together with one-sample *t*-tests within each Group × Direction condition to test whether ΔIHI differed from zero.

To examine whether individual differences in TMS-derived measures were related to behavioral improvements, we computed correlations between ΔMEP (Experiment 2) or ΔIHI (Experiment 3) and both training-related (online; Late − Early) and consolidation-related (offline; 6 h − Late) changes in coordination accuracy and variability. Spearman’s *ρ* was used as the primary statistic given the relatively small sample size per group; Pearson correlations were computed in parallel and yielded equivalent conclusions throughout. Correlations were performed separately for each Group × Hemisphere combination using corr with the ‘Type’, ‘Spearman’ and ‘Type’, ‘Pearson’ options.

## Results

### Experiment 1: Acquisition and consolidation of the 90° relative phase pattern

Experiment 1 examined whether the novel 90° relative phase pattern could be acquired through practice in the three coordination contexts—bimanual, unimanual-left, and unimanual-right—and whether the acquired performance was retained 6-h after training. These shared signatures in the present context equate the behavioral task across experiments and provide a balanced comparison of the TMS-derived measures in experiments 2 and 3.

#### Coordination accuracy

Across training, coordination became increasingly more accurate in all three groups, as reflected by increases in BW30 (Figure 2A–B) and by a tightening of the relative phase distribution around 90° (Figure 2J–R). Accuracy was higher again after the 6-h interval, a pattern consistent with consolidation-related offline improvement (Figure 2A–B, J–R).

**Figure 2.**
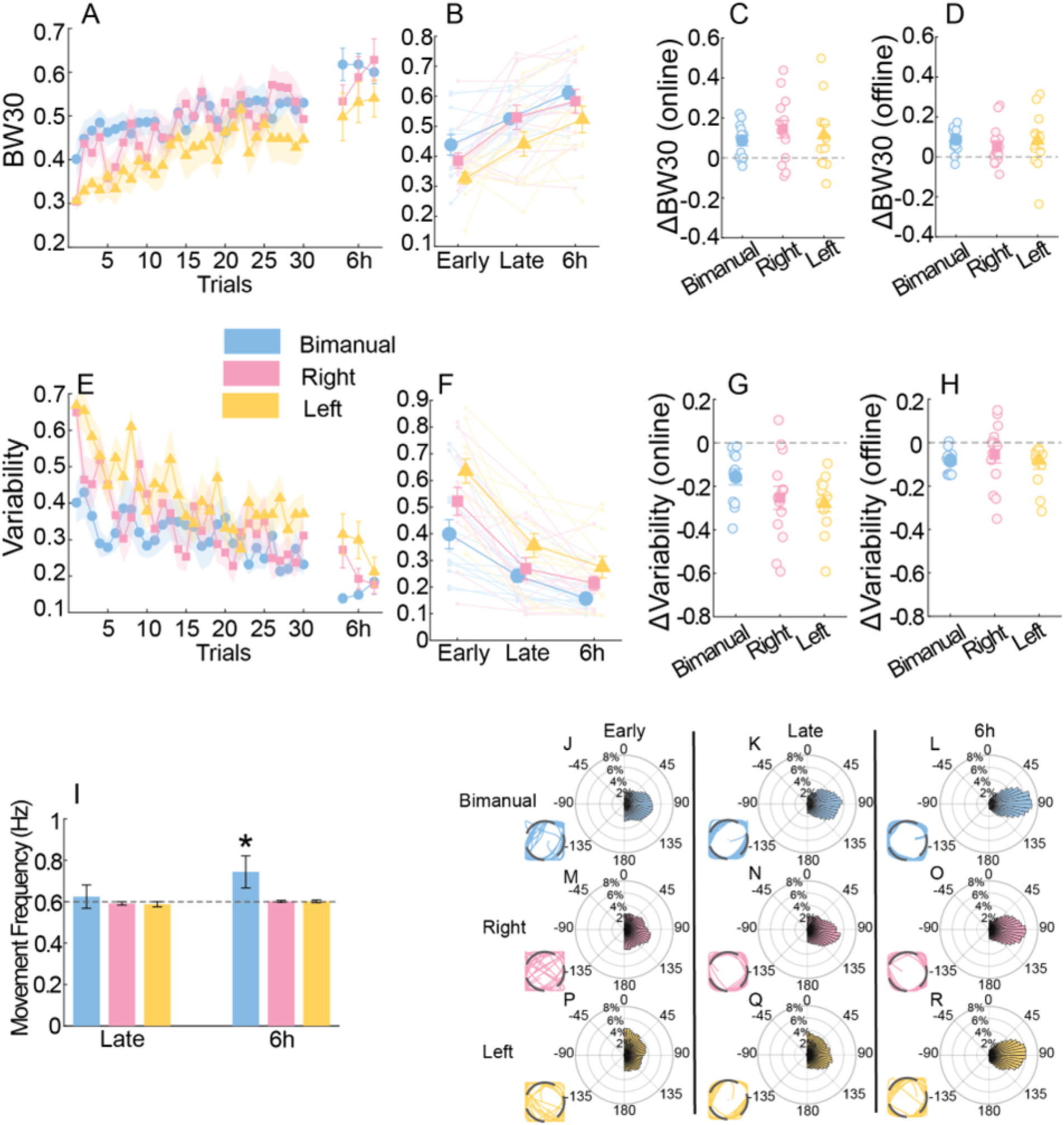
Behavior for Experiment 1. A) Proportion of time (BW30) within ±30° of the target 90° pattern during practice and the 6-h retention test. Higher values indicate better coordination accuracy. B) Mean BW30 during early practice, late practice, and the 6-hour retention test. C) Online improvement in coordination accuracy, calculated as the difference between late- and early-practice BW30. D) Offline improvement over the 6-hour post-training interval, calculated as the difference between 6-h retention and late-practice BW30. E) Coordination (*R*)variability during initial practice and the 6-h retention test. F) Mean coordination variability during early practice, late practice, and the 6-h retention test. G) Online improvement in coordination variability, with the negative value indicating increased coordination stability. H) Offline improvement in coordination variability. I) Movement frequency (Hz) during late practice and the 6-h retention test. The dashed line indicates the 0.6-Hz movement frequency imposed by the virtual partner during unimanual coordination. J – R) Polar histograms of continuous relative phase for early practice, late practice, and the 6-h retention test. The distributions progressively narrow and more tightly centered around 90°, reflecting improved performance. Representative real-time feedback trajectories are shown for each stage of practice and retention across the three training groups. Asterisks indicate statistical significance at p < .05.

The linear mixed-effects model applied to the BW30 data revealed a significant main effect of Time, *F*(2, 105) = 32.79, *p* < .001, η_p_^2^ = .384, indicating progressive improvement in coordination accuracy over time. The main effect of Group was also significant, *F*(2, 105) = 4.11, *p* = .019, η_p_^2^ = .073. The Group × Time interaction was not significant, *F*(4, 105) = 0.35, *p* = .841, indicating that the time course of improvement was comparable across groups (Figure 2B–D). Planned pairwise comparisons for the main effect of Time indicated that coordination accuracy at Late training was better than at Early training (*b* = 0.12, *SE* = 0.02, *t*(105) = 5.34, *p* < .001), confirming training-related online improvement. Coordination accuracy at 6-h was greater than at Late training (*b* = 0.07, *SE* = 0.02, *t*(105) = 3.44, *p* < .001) and Early training (*b* = 0.19, *SE* = 0.02, *t*(105) = 8.78, *p* < .001), consistent with consolidation-related offline improvement. Planned pairwise comparisons for the main effect of Group showed that the bimanual group was more accurate than the unimanual-left group (*b* = 0.09, *SE* = 0.03, *t*(111) = 2.76, *p* = .01), and that the unimanual-right group was also more accurate than the unimanual-left group (*b* = 0.07, *SE* = 0.03, *t*(105) = 2.09, *p* = .03). Accuracy did not differ between the bimanual and unimanual-right groups (*b* = 0.03, *SE* = 0.03, *t*(105) = 0.77, *p* = .44).

#### Coordination variability

Coordination became progressively more stable around 90° across training, and stability improved further after the 6-h interval (Figure 2E–H, J–R). The linear mixed-effects model applied to the adjusted *R* values revealed a significant main effect of Time, *F*(2, 105) = 66.77, *p* < .001, η_p_^2^ = .560, and a significant main effect of Group, *F*(2, 105) = 6.24, *p* = .003, η_p_^2^ = .106. The Group × Time interaction was not significant, *F*(4, 105) = 1.22, *p* = .31. Planned pairwise comparisons for the main effect of Time indicated that coordination variability at Late training was smaller than at Early training (*b* = 0.23, *SE* = 0.02, *t*(105) = 9.65, *p* < .001), confirming training-related online improvement. Variability at 6-h was smaller than at Late training (*b* = 0.07, *SE* = 0.02, *t*(105) = 3.10, *p* = .002) and Early training (*b* = 0.30, *SE* = 0.02, *t*(105) = 12.74, *p* < .001), consistent with consolidation-related offline improvement. Planned pairwise comparisons for the main effect of Group indicated that variability in the unimanual-left group was greater than in the bimanual group (*b* = −0.16, *SE* = 0.04, *t*(105) = −3.52, *p* < .001) and greater than in the unimanual-right group (*b* = 0.09, *SE* = 0.04, *t*(105) = −2.04, *p* = .04). Variability did not differ between the bimanual and unimanual-right groups (*b* = −0.07, *SE* = 0.04, *t*(105) = −1.62, *p* = .11).

#### Movement frequency

The linear mixed-effects model fit to the *f_dom_* data revealed that the main effect of Group was not significant, *F*(2, 70) = 2.06, *p* = .13, η_p_^2^ = .06. The analysis did reveal a significant main effect of Time, *F*(1, 70) = 20.80, *p* < .001, η_p_^2^ = .23, and a significant Group × Time interaction, *F*(2, 70) = 11.33, *p* < .001, η_p_^2^ = .25 (Figure 2E). Planned pairwise comparisons of the Group × Time interaction indicated that the bimanual group moved at a faster movement frequency at the 6-h retention test than at Late training (*b* = 0.12, *SE* = 0.02, *t*(70) = 6.40, *p* < .001). In contrast, movement frequency remained comparable between Late training and the 6-h retention test in the unimanual-left (*b* = 0.01, *SE* = 0.02, *t*(70) = 0.74, *p* = .46) and unimanual-right groups (*b* = 0.01, *SE* = 0.02, *t*(70) = 0.61, *p* = .54). Participants in the unimanual-left and unimanual-right groups had to synchronize their movements with a virtual partner moving at 0.6 Hz. Separate one-sample *t* tests confirmed that movement frequency in these two groups did not differ from 0.6 Hz (all *p*s > .30).

#### Summary Experiment 1

Overall, training facilitated learning the novel 90° relative phase pattern with minimal differences in online and offline learning rates between the three training groups. The changes in coordination accuracy and coordination variability demonstrate the development of a stable attractor at the targeted relative phase pattern of 90°. Interestingly, movement frequency increased in the bimanual group between the end of training and retention. Typically, an increase in movement frequency destabilizes coordination patterns, yet the 90° pattern remained stable across the retention interval and further the stabilization of an attractor at the 90° relative phase. Differences in movement frequency are therefore unlikely to explain the group patterns observed in coordination performance. The lower accuracy and slightly higher variability observed in the unimanual-left group may reflect a hand-dominance effect, given that the nondominant left hand is associated with greater variability and slower movement speed in rhythmic tapping tasks (Guiard, 1987; Peters, 1994). Importantly, this baseline difference does not undermine the central observation that coordination accuracy in the unimanual-left group improved over time. Given the comparable temporal dynamics of performance changes in training and after the offline interval, we next asked whether the three forms of practice are supported by similar or distinct neural mechanisms in Experiment 2.

### Experiment 2: Modulation of bilateral corticospinal excitability by unimanual and bimanual training

In this experiment, the behavioral protocol of Experiment 1 was repeated and single-pulse TMS over left M1 and right M1 were administered before and after training to measure training-related changes in corticospinal excitability (Figure 1F).

#### Coordination accuracy

Coordination accuracy increased in all groups across training (Figure 3A–B), indicating learning-related improvement in producing the 90° pattern. The analysis of the BW30 data revealed a significant main effect of Time, *F*(2, 105) = 55.79, *p* < .001, η_p_^2^ = .52, and a marginally significant main effect of Group, *F*(2, 105) = 3.05, *p* = .05, η_p_^2^ = .055. Planned pairwise comparisons for the main effect of Time indicated that coordination accuracy was greater during Late training than during Early training (Early – Late: *b* = −0.12, *SE* = 0.02, *t*(105) = −5.84, *p* < .001), indicating training-related online improvement. Moreover, accuracy was greater after the 6-h retention interval compared to Early training (Early – 6 h: *b* = −0.21, *SE* = 0.02, *t*(105) = −10.56, *p* < .001), and Late training (Late – 6 h: *b* = −0.09, *SE* = 0.02, *t*(105) = −4.67, *p* < .001), consistent with consolidation-related offline improvement. Planned pairwise comparisons for the main effect of Group indicated that the bimanual group was marginally more accurate than the unimanual-left group (bimanual – unimanual-left: *b* = 0.08, *SE* = 0.04, *t*(105) = 1.92, *p* = .06) and significantly more accurate than the unimanual-right group (bimanual – unimanual-right: *b* = 0.09, *SE* = 0.04, *t*(105) = 2.30, *p* = .02). Coordination accuracy was comparable between the unimanual-left and unimanual-right groups (unimanual-right – unimanual-left: *b* = −0.01, *SE* = 0.04, *t*(105) = −0.33, *p* = .74). The Group × Time interaction was not significant, *F*(4, 105) = 0.88, *p* = .48, η_p_^2^ = .032 (Figure 3C–D).

**Figure 3.**
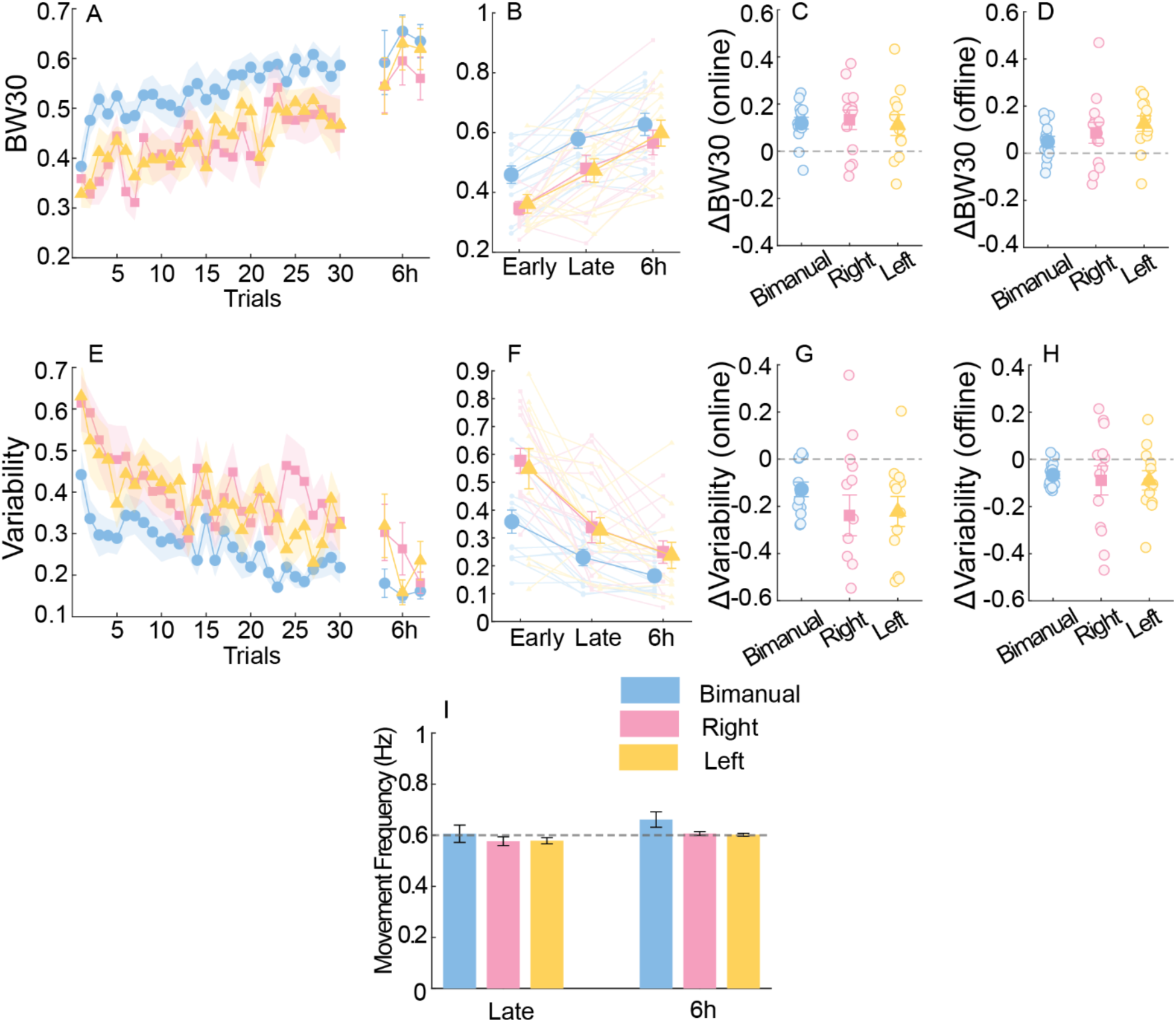
Behavior for Experiment 2. A) Proportion of time (BW30) within ±30° of the target 90° phase during practice and at 6-hour retention test. Higher values indicate greater coordination accuracy. B) Mean BW30 during early practice, late practice, and at the 6-hour retention test. C) Online improvement in coordination accuracy, calculated as the difference between late- and early-practice (BW30). D) Offline improvement calculated as the difference between 6-hour retention and late-practice (BW30). E) Coordination variability during practice and the 6-hour retention test. F) Mean coordination variability during early practice, late practice, and the 6-hour retention test. The corresponding values were calculated using the same procedure described above for the accuracy measures. G) Online improvement in coordination variability, with the negative value indicating the improvement in stability. H) Offline improvement in coordination variability. I) Movement frequency (Hz) during late practice and the 6-hour retention test. The dashed line indicates the 0.6-Hz movement frequency imposed by the virtual partner during unimanual coordination.

#### Coordination variability

Coordination variability decreased gradually in all three groups across training and across the 6-h interval (Figure 3E–H). The analysis of the variability data (*1-R*) revealed a significant main effect of Time, *F*(2, 105) = 42.62, *p* < .001, η_p_^2^ = .446, and a significant main effect of Group, *F*(2, 105) = 5.95, *p* = .004, η_p_^2^ = .102. Planned pairwise comparisons for the main effect of Time indicated that variability at Late training was lower than at Early training (*b* = 0.20, *SE* = 0.03, *t*(105) = 6.32, *p* < .001), indicating training-related online improvement. Coordination variability at the 6-h retention test was lower than at Early training (*b* = 0.28, *SE* = 0.03, *t*(105) = 8.94, *p*< .001) and Late training (*b* = 0.08, *SE* = 0.03, *t*(105) = 2.63, *p* = .010), consistent with consolidation-related offline improvement. Planned pairwise comparisons for the main effect of Group indicated that variability was lower in the bimanual group than in both the unimanual-left group (*b* = −0.12, *SE* = 0.04, *t*(105) = −2.72, *p* = .008) and the unimanual-right group (*b* = −0.14, *SE* = 0.04, *t*(105) = −3.18, *p* = .002). Variability did not differ between the unimanual-left and unimanual-right groups (*b* = 0.02, *SE* = 0.04, *t*(105) = 0.40, *p* = .690). The Group × Time interaction was not significant, *F*(4, 105) = 1.06, *p* = .380.

#### Movement frequency

The fit of the linear mixed-effects model to the *f_dom_* data revealed a significant main effect of Time, *F*(1, 70) = 10.10, *p* = .002, η_p_^2^ = .126, indicating that movement frequency was higher at 6-h retention than at Late training (Figure 3I). Neither the main effect of Group, *F*(2, 70) = 1.93, *p* = .153, nor the Group × Time interaction, *F*(2, 70) = 0.75, *p* = .476, was significant. Movement frequency in the unimanual-left and unimanual-right groups did not differ from the virtual partner’s frequency of 0.6 Hz at either Late training or at the 6-h retention test (all *p*s > .11), indicating that participants’ movements were well synchronized with the virtual partner (Figure 3I).

#### Bilateral corticospinal excitability

Cortical excitability change as a function of practice was assessed with a linear mixed-effects model analysis of the ΔMEP data with positive values indicating a training-related increase. The intercept was significant, *F*(1, 70) = 25.00, *p* < .001, indicating an overall increase in MEP amplitude from pre- to post-training across groups and hemispheres. The main effect of Hemisphere was marginal, *F*(1, 70) = 3.47, *p* = .067, η_p_^2^ = .047, while neither the main effect of Group, *F*(2, 70) = 0.79, *p* = .457, η_p_^2^ = .022, nor the Group × Hemisphere interaction, *F*(2, 70) = 0.02, *p* = .980, η_p_^2^ = .001, reached significance. Exploring the marginal hemisphere effect with pairwise comparisons indicated a numerically larger excitability increase at the dominant left M1 than at the right M1 (*b* = −0.16, *SE* = 0.09, *t*(70) = −1.86, *p* = .067; Figure 4C–F), (Figure 4A–B, D–E).

**Figure 4.**
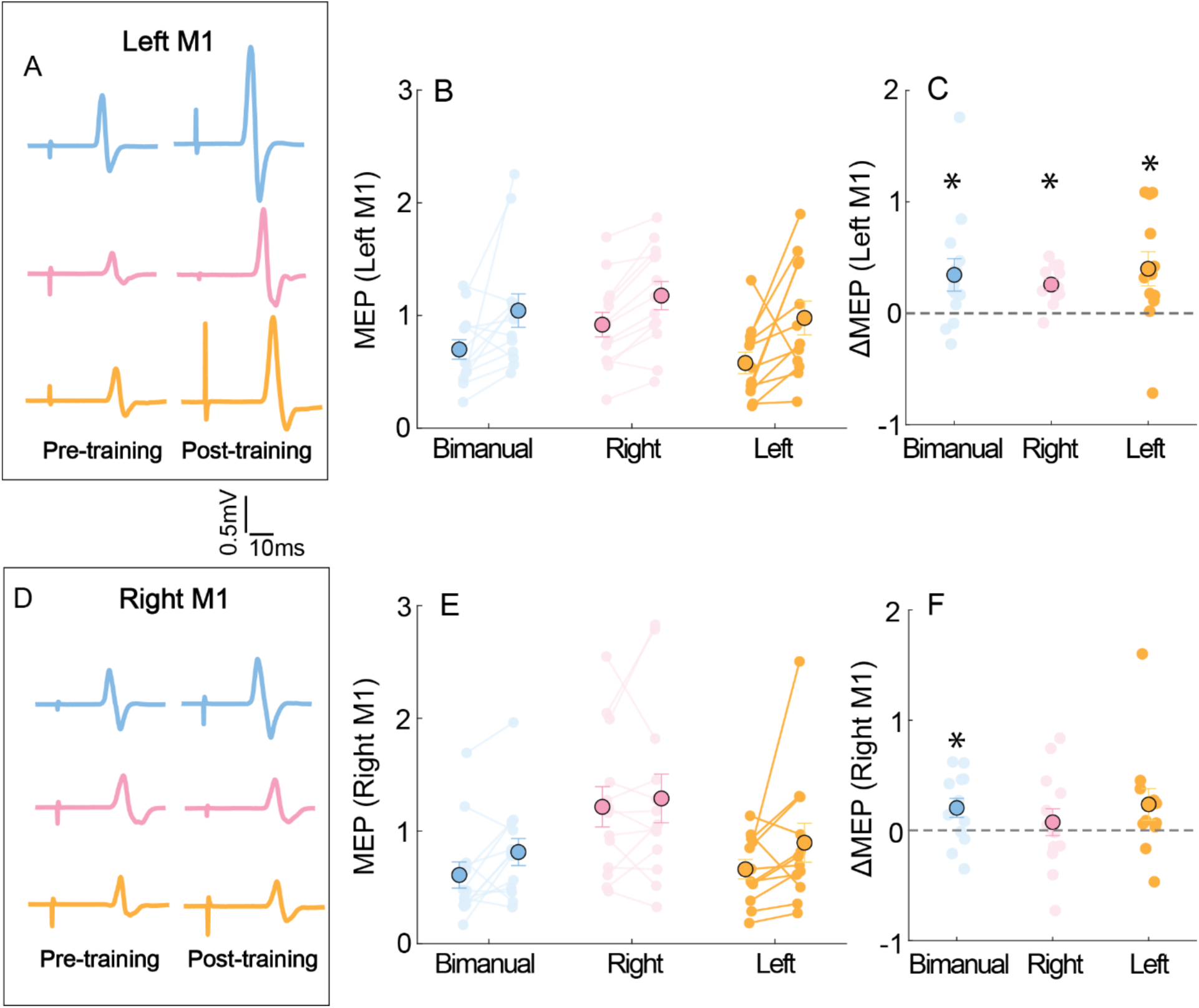
A) For each group, a representative motor evoked (MEP) potential measured from left M1 during pre- and post-training. B) Mean MEPs from left M1 during pre- and post-training across three groups. Scatter dots represent individual subjects MEPs. C) The change in motor evoked potential for all three groups in left M1 is reflected by the difference between pre- and post-training. Asterisks reflect the significant difference relative to zero by one-sample t test. D) For each group, a representative MEP measured from right M1 during pre- and post-training. E) Mean MEP from the right M1 during pre- and post-training across three groups. Scatter dots represent individual subjects MEPs. F) The change in motor evoked potential in the right M1 for the bimanual group is reflected by the difference between pre- and post-training. Asterisk indicates statistical significance at p < .05 from one-sample t test against zero.

We additionally conducted one-sample *t* tests to determine whether the MEP change in each Group × Hemisphere condition differed from zero. Bimanual practice was associated with a significant increase in MEP amplitude at both the left M1 (*M* = 0.20 ± 0.31, *t*(12) = 2.37, *p* = .04) and the right M1 (*M* = 0.34 ± 0.53, *t*(12) = 2.36, *p* = .04). For the unimanual-right group, a significant increase was observed only at left M1 (*M* = 0.25 ± 0.16, *t*(12) = 5.70, *p* < .001), whereas right M1 was unchanged (*M* = 0.07 ± 0.44, *t*(12) = 0.61, *p* = .55). For the unimanual-left group, MEP amplitude increased significantly at left M1 (*M* = 0.40 ± 0.53, *t*(11) = 2.62, *p* = .02) but not at right M1 (*M* = 0.24 ± 0.50, *t*(11) = 1.64, *p* = .13). These follow-up tests indicate that MEP increases were reliably greater than zero in the left M1 in all three groups, whereas increases in the right M1 reached significance only after bimanual practice. (Figure 5C, F).

**Figure 5.**
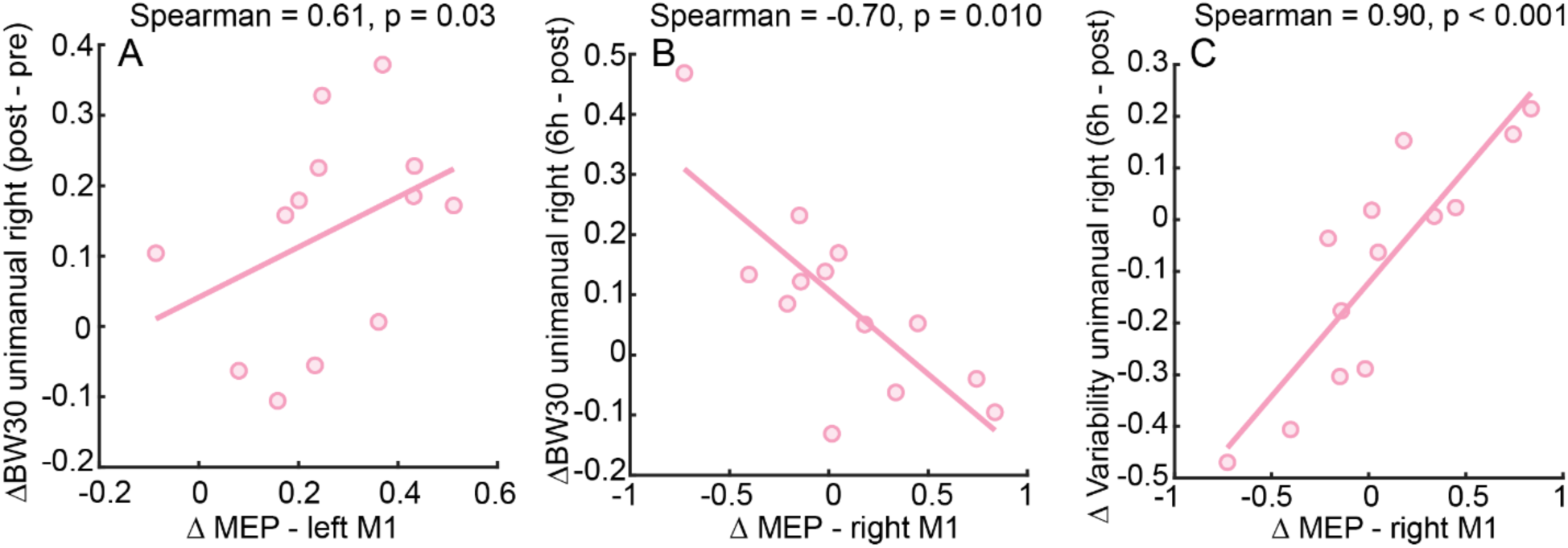
Correlational analyses between changes in motor-evoked potential (MEP) amplitude and behavioral performance in Experiment 2. **A)** Correlation between changes in left M1 MEP amplitude and practice-related changes in coordination accuracy in the unimanual-right group. **B)** Correlation between changes in right M1 MEP amplitude and changes in coordination accuracy from post-training to the 6-hour retention test. **C)** Correlation between changes in right M1 MEP amplitude and changes in coordination variability from post-training to the 6-hour retention test.

Previous work has implicated M1 in both the acquisition and the consolidation of motor memories, we examined whether individual differences in corticospinal excitability change were related to behavioral improvement. We correlated MEP changes with training-related (online) and consolidation-related (offline) changes in coordination accuracy and variability. In the unimanual-right group, MEP change at the dominant left M1 was positively associated with online improvement in coordination accuracy (Spearman’s ρ = 0.61, *p* = .03; Figure 5A). Also in the unimanual-right group, MEP change at the nondominant right M1 was negatively associated with offline change in coordination accuracy (Spearman’s ρ = −0.70, *p* = .01; Figure 5B) and positively associated with offline change in coordination variability, for which smaller values indicate more stable coordination (Spearman’s ρ = 0.90, *p* < .01; Figure 5C). Greater increases in right M1 excitability after right-hand practice were thus associated with smaller offline gains, as reflected by lower accuracy and greater variability at 6-h retention. No other correlations were significance (see. Tables 1–2 in Supplementary materials).

#### Summary Experiment 2

The performance changes that emerged were very consistent with experiment 1, especially the online and offline changes found in the groups during training. Experiment 1 suggested the possibility of a handedness effect; however, the second experiment did not reveal the same set of significant effects to support a similar handedness outcome across the two experiments. TMS revealed an overall increase in M1 excitability in both hemispheres in post-practice compared to pre-practice. There were no group related effects or hemispheric differences identified in the main analysis, follow up t-tests did find that excitability increased in both hemispheres for the bimanual group, and only in the contralateral hemispheres for the unimanual-right and unimanual-left hand trained groups. Correlational analysis did reveal a possible handedness asymmetry in the TMS data, with significant correlations found between cortical excitability in left M1 and right M1 as a function of coordination accuracy (left M1) and stability (right M1) changes with practice. These associations raise the possibility that stabilization of the 90° attractor during unimanual virtual-partner practice depends on excitability changes within each M1 and how activity is balanced between the hemispheres. This modest between-hemisphere difference in corticospinal excitability change hint that bimanual and unimanual coordination rely on bilateral neural substrates supported by interhemispheric communication. This was explored in Experiment 3 with the use of a TMS IHI protocol.

### Experiment 3: Modulation of Interhemispheric inhibition by unimanual and bimanual training

This experiment tested directly whether unimanual and bimanual practice differentially modulated IHI by using a dual-site paired-pulse TMS protocol (Figure 1G). This protocol allowed us to examine whether the behavioral improvements emerging online during training and over offline interval were accompanied by practice-specific changes in inhibitory communication between the two motor cortices.

#### Coordination accuracy

Coordination performance improved progressively across training and the 6-h retention interval in all three groups (Figure 6A–B). The analysis of the BW30 data revealed a significant main effect of Time, *F*(2, 99) = 136.66, *p* < .001, η_p_^2^ = .734. Neither the main effect of Group, *F*(2, 99) = 0.42, *p* = .660, nor the Group × Time interaction, *F*(4, 99) = 0.62, *p* = .646, was significant (Figure 6C–D). Planned pairwise comparisons for the main effect of Time confirmed that coordination accuracy at Late training was better than at Early training (Late – Early: *b* = 0.23, *SE* = 0.018, *t*(99) = 12.35, *p* < .001), confirming on-line related improvement. Coordination accuracy at the 6-h retention test was better than at Early training (6 h – Early: *b* = 0.29, *SE* = 0.018, *t*(99) = 15.69, *p* < .001) and better than Late training (6 h – Late: *b* = 0.061, *SE* = 0.018, *t*(99) = 3.35, *p* = .001), confirming consolidation-related offline improvement.

**Figure 6.**
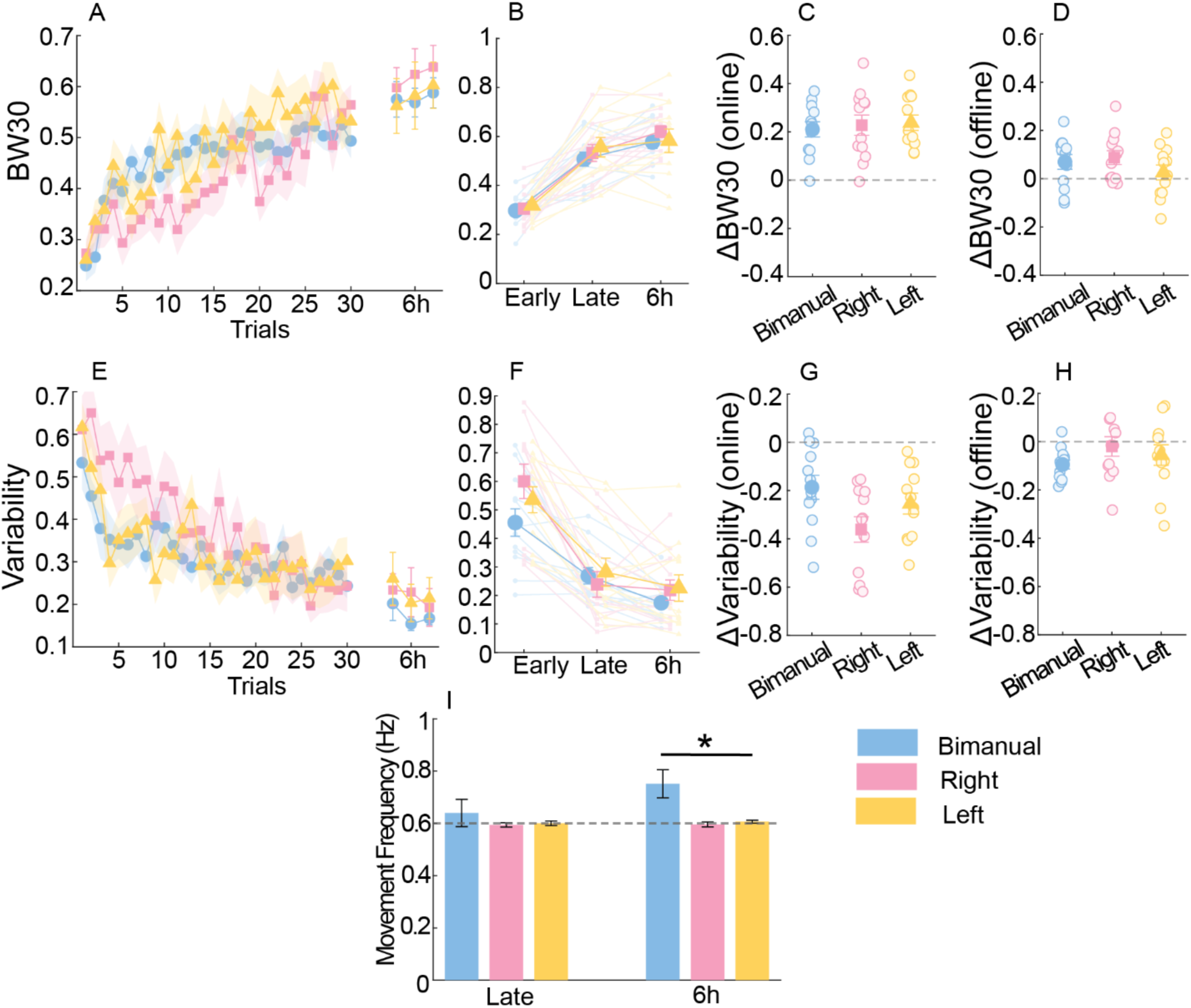
Behavior for Experiment 3. A) Learning curves reflected by BW30 across three groups. Higher values indicate greater coordination accuracy. B) Mean BW30 during early practice, late practice, and the 6-hour retention test. C) Online improvement in coordination accuracy (BW30). D) Offline improvement over the 6-hour post-training interval, calculated as the difference between 6-hour retention and late-practice BW30. E) Coordination variability during initial practice and the 6-hour retention test. F) Mean coordination variability during early practice, late practice, and the 6-hour retention test. G) Online improvement in coordination variability, with the negative value indicating the improvement in stability. H) Offline improvement in coordination variability. I) Movement frequency (Hz) during late practice and the 6-hour retention test. Asterisk indicates statistical significance at p < .05.

#### Coordination variability

Coordination variability showed the same overall pattern as in the previous two experiments with variability decreasing progressively across training and decreasing further across the 6-h retention interval in the three groups (Figure 6E–F). The linear mixed-effects model revealed a significant main effect of Time, *F*(2, 99) = 90.06, *p* < .001, η_p_^2^ = .645, and a nonsignificant main effect of Group, *F*(2, 99) = 0.79, *p* = .455. The Group × Time interaction did not reach significance, *F*(4, 99) = 1.98, *p* = .103 (Figure 6G–H). Planned pairwise comparisons for the main effect of Time confirmed that variability at Late training was smaller than at Early training (Early – Late: *b* = 0.27, *SE* = 0.026, *t*(99) = 10.36, *p* < .001), indicating online improvement. Variability at the 6-h retention test was smaller than at Early training (Early – 6 h: *b* = 0.32, *SE* = 0.026, *t*(99) = 12.57, *p* < .001) and smaller than at Late training (Late – 6 h: *b* = 0.057, *SE* = 0.026, *t*(99) = 2.20, *p* = .030), indicating offline improvement in performance.

#### Movement frequency

The linear mixed-effects model analysis of the *f_dom_* frequency data revealed a significant main effect of Time, *F*(1, 66) = 11.62, *p* = .001, η_p_^2^ = .15, a significant main effect of Group, *F*(2, 66) = 3.90, *p* = .025, η_p_^2^ = 0.11, and a significant Group × Time interaction, *F*(2, 66) = 9.49, *p* < .001, η_p_^2^ = 0.22, indicating that the pattern of frequency change over time differed across groups (Figure 6I). To decompose the Group × Time interaction, the model was re-estimated with different reference conditions. The bimanual group showed a higher movement frequency at 6-h retention than at Late training (6 h – Late: *b* = 0.11, *SE* = 0.020, *t*(66) = 5.52, *p* < .001), replicating the pattern observed in Experiment 1. In contrast, movement frequency remained comparable between Late training and 6-h retention test in the unimanual-right group (6-h – Late: *b* = 0.0015, *SE* = 0.020, *t*(66) = 0.076, *p* = .940) and unimanual-left group (6-h – Late: *b* = 0.0015, *SE* = 0.020, *t*(66) = 0.076, *p* = .940). Movement frequency at the 6-h retention test was lower in the unimanual-right group (*b* = −0.156, *SE* = 0.042, *t*(66) = −3.67, *p* < .001) and unimanual-left group compared to the bimanual group (*b* = −0.145, *SE* = 0.042, *t*(66) = −3.41, *p* = .001). Consistent with the requirement that the unimanual groups synchronize with the virtual partner, one-sample *t* tests confirmed that movement frequency in the unimanual-right group did not differ from 0.6 Hz at Late training, *t*(11) = −0.73, *p* = .481, or at 6-h retention, *t*(11) = −0.46, *p* = .653, and that movement frequency in the unimanual-left group did not differ from 0.6 Hz at Late training, *t*(11) = 0.04, *p*= .972, or at 6-h retention, *t*(11) = 1.15, *p* = .275.

#### Interhemispheric inhibition

To examine whether unimanual and bimanual practice differentially modulated interhemispheric communication, a linear mixed-effects model was fit to the ΔIHI change scores. The analysis revealed a significant Group × Direction interaction, *F*(2, 66) = 3.17, *p* = .048, indicating that the pattern of IHI change differed between groups as a function of the direction probed. Neither the main effect of Group, *F*(2, 66) = 0.61, *p* = .548, nor the main effect of Direction, *F*(1, 66) = 3.23, *p* = .077, was significant (Figure 7B–E). To decompose the interaction, we conducted planned comparisons by re-estimating the model with different reference conditions and performed one-sample *t* tests within each Group × Direction condition to determine whether the change in IHI differed from zero.

**Figure 7.**
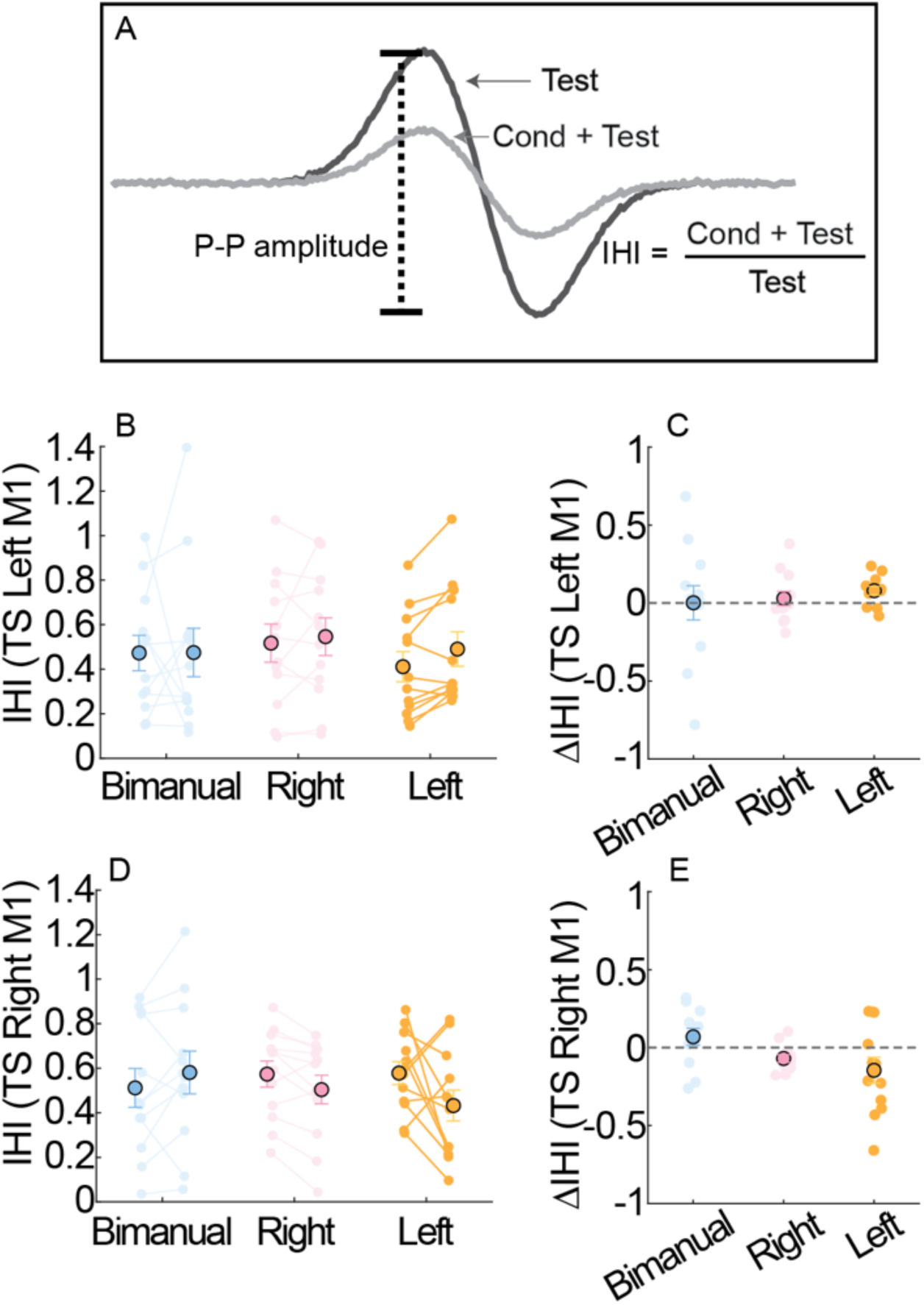
A) The illustration of interhemispheric inhibition (IHI) index. B) Mean IHI index from testing the left M1 during pre- and post-training across three groups. Scatter dots represent individual’s motor evoked potential. C) The change in IHI index from testing the left M1 reflected by the difference between pre- and post-training. D) Mean IHI index from testing the right M1 during pre- and post-training across three groups. Scatter dots represent individual’s motor evoked potential. F) The change in IHI index from testing the right M1 reflected by the difference between pre- and post-training.

For the bimanual group, the planned comparison between the two testing directions revealed no significant difference, *b* = 0.07, *SE* = 0.08, *t*(66) = 0.82, *p* = .42 (Figure 7C, E), indicating that practice-induced changes in inhibition directed toward the left and right M1 were comparable. Relative to pre-training, IHI in the bimanual group was unchanged when the right M1 was tested (*t*(11) = 1.28, *p* = .23) and when the left M1 was tested (*t*(11) = 0.02, *p* = .98). Between groups, the change in the inhibition exerted by left M1 on right M1 differed between the unimanual-left and bimanual groups (*b* = −0.21, *SE* = 0.09, *t*(66) = −2.44, *p* = .02) but not between the unimanual-right and bimanual groups (*b* = −0.14, *SE* = 0.09, *t*(66) = −1.57, *p* = .12) or between the unimanual-left and unimanual-right groups (*b* = 0.08, *SE* = 0.09, *t*(66) = 0.86, *p* = .39).

In the unimanual-left group, the change in IHI differed between the two testing directions (*b* = 0.23, *SE* = 0.08, *t*(66) = 2.73, *p* = .01; Figure 7C, E). When left M1 was tested, the change was positive, reflecting a reduction in the inhibition that right M1 (contralateral to the trained hand) exerted on left M1. One-sample tests confirmed this reduction (*t*(11) = 2.90, *p* = .01), whereas the inhibition received by right M1 was unchanged (*t*(11) = −1.76, *p* = .11), showing only a numerical trend toward stronger inhibition.

In the unimanual-right group, the change in IHI did not differ between the two testing directions (*b* = 0.10, *SE* = 0.08, *t*(66) = 1.20, *p* = .23). Relative to pre-training, the inhibition that left M1 exerted on right M1 increased significantly (*t*(11) = −2.78, *p* = .02), whereas the inhibition received by the left M1 was unchanged (*t*(11) = 0.63, *p* = .54).

Next, we examined whether individual differences in IHI change were related to behavioral improvement. Spearman correlations were computed as the primary analysis given the small sample size (n = 12 per group), with Pearson correlations reported for completeness. We tested associations between the change in IHI in each Group × Direction condition and both training-related (online) and consolidation-related (offline) changes in coordination accuracy and variability. No significant correlations emerged between the change in IHI and online behavioral gains (all Spearman *p*s > .10) or between the change in IHI and offline gains (all Spearman *p*s > .22); Pearson correlations yielded the same conclusions (see Table 3 – 4 in supplementary materials). At the level of individual differences, the magnitude of training-related change in IHI was therefore not systematically related to the extent of behavioral improvement in Experiment 3. *Summary Experiment 3*

The behavioral learning profile observed in Experiments 1 and 2 was replicated, confirming that comparable online and offline improvements in coordination performance occurred under all three practice conditions. Critically, dual-site paired-pulse TMS revealed a significant Group × Direction interaction in training-related IHI change. Right-hand unimanual practice strengthened the inhibition that the left M1 exerted on the right M1 while leaving the opposite pathway unchanged. Left-hand unimanual practice showed the complementary pattern: the inhibition that the right M1 exerted on the left M1 was reduced. Bimanual practice, in contrast, produced no reliable change in either direction. To our knowledge, these are the first TMS data indicating that learning the same coordination pattern under unimanual and bimanual training contexts are accompanied by different patterns of interhemispheric plasticity, with the direction of change in each unimanual group consistent with the asymmetric inhibitory organization of the motor system in right-handed individuals.

## Discussion

To our knowledge, this is the first study to determine whether learning the same novel 90° relative phase pattern through bimanual and unimanual coordination engages shared or effector-dependent neurophysiological processes across the two M1s. Across three experiments, all coordination conditions showed progressive improvements in accuracy and stability during practice, followed by further improvement over the 6-hour post-training interval, indicating comparable online and offline behavioral changes across effector configurations. Against this common behavioral background, the neurophysiological findings revealed a dissociation between corticospinal excitability and interhemispheric interactions. Training produced an overall increase in corticospinal excitability across both M1s that did not differ across coordination conditions, whereas interhemispheric inhibition varied with the effector configuration and, across both unimanual conditions, showed a pattern consistent with preserving the functional influence of the dominant left M1.

### The acquisition and consolidation of a 90-degree coordination pattern

Consistent with previous findings, the present study is the first to show across three independent experiments that Lissajous feedback reliably supports rapid acquisition of a novel 90° relative-phase pattern as a stable attractor in unimanual and bimanual tasks, with a comparable pattern of improvement observed in each experiment (Buchanan & Wang, 2012; Kovacs et al., 2009b, 2009c, 2010a, 2010b; Wang, Weinrich, et al., 2024). A key distinction between unimanual and bimanual coordination, however, lies in the effector configuration through which the action outcome is generated. During bimanual coordination, cursor motion reflects the continuous integration of movements produced by the left and right hands, such that coordination is supported by both shared visual information and interactions between bilateral motor systems (Carson, 2005; Swinnen, 2002). In contrast, during unimanual coordination, movement produced by a single hand must be synchronized with an externally generated motion along the orthogonal dimension. This difference in effector configuration may alter the strength and stability of the underlying coordination coupling. Indeed, previous work has shown that unimanual coordination can be less stable than bimanual coordination, potentially reflecting the weaker, unidirectional coupling between the active hand and an externally generated partner compared with the reciprocal coupling available between two actively controlled limbs (Kelso et al., 2009; Snapp-Childs et al., 2011, 2015). The present results showed a similar tendency, with modest advantages in bimanual accuracy and stability observed in Experiments 1 and 2, although these differences were not consistently present across all three experiments. Critically, however, the time course of practice-related improvement was comparable across coordination conditions, suggesting that differences in coupling strength did not substantially constrain the capacity to acquire the 90° pattern.

Although Lissajous feedback enables rapid stabilization of a novel coordination pattern, performance can deteriorate when the integrated visual feedback is removed, leading these practice effects to be viewed primarily as a short-term adaptive process strongly dependent on concurrent visual information (Kovacs et al., 2009a). Evidence of post-training improvement, however, suggests that learning supported by Lissajous feedback is not entirely transient (I. Park et al., 2020). In the present study, coordination accuracy and stability improved further across the 6-hour post-training interval in all three groups, indicating reliable offline improvement after practice had ended. A similar pattern has been reported for bimanual coordination learning, with Park and colleagues (2020) demonstrating further improvements in accuracy and stability following 30-minute and 6-hour post-training intervals. McCulloch and colleagues (2021) subsequently characterized the time-dependent stabilization of bimanual coordination memory using a retrograde interference paradigm (McCulloch et al., 2021; Inchon Park et al., 2020). Learning a competing coordination pattern 2 hours after initial practice interfered with the previously acquired memory, whereas introducing the competing pattern after a 6-hour interval did not impair subsequent performance of the original pattern. These findings parallel the broader motor-memory literature showing time-dependent stabilization or enhancement following unimanual motor sequence learning (Robertson et al., 2004; Walker et al., 2002) and sensorimotor adaptation (Brashers-Krug et al., 1996; Krakauer & Shadmehr, 2006; Wang, Huynh, et al., 2024). The present findings therefore suggest that memories acquired through unimanual rhythmic coordination may undergo a similar post-training stabilization process. Nevertheless, offline improvement alone does not definitively establish consolidation. Future studies using retrograde interference at different post-training intervals will be necessary to determine whether rhythmic unimanually acquired coordination memories become progressively resistant to interference and whether their temporal dynamics parallel those established for bimanual coordination.

### Bilateral corticospinal excitability changes across coordination conditions

The consistent accuracy and stability changes observed for the unimanual and bimanual tasks provide a foundation for interpreting the subsequent TMS findings. The neurophysiological changes measured across the two M1s can be considered in relation to acquisition of a common stable attractor (90° pattern) rather than learning of entirely different task-specific actions. Previous TMS studies examining corticospinal involvement in bimanual coordination have primarily assessed the M1 contralateral to the dominant hand. In general, these studies revealed that practicing a 90° relative phase pattern bimanually increased corticospinal excitability in the left (dominant) hemisphere (Nomura et al., 2016; Inchon Park et al., 2020). Extending this previous work, the present study assessed both M1s and revealed an overall training-related increase in corticospinal excitability in M1 across hemispheres, regardless of whether training involved bimanual coordination or unimanual coordination with either hand. Although the increase was numerically greater in the dominant left M1, the hemispheric difference did not reach statistical significance, and the pattern of excitability change did not differ across training conditions. These findings therefore suggest that acquisition of the 90° coordination pattern is accompanied by a broadly bilateral increase in corticospinal excitability rather than a response confined to the M1 contralateral to the actively trained hand.

In the current study, unimanual coordination performed with the left and right hands served as complementary conditions for identifying hemisphere-specific neurophysiological changes associated with acquiring the same spatiotemporal coordination pattern. A strictly effector-dependent account would predict that practice with one hand should primarily modulate the contralateral M1, whereas increases in corticospinal excitability within the M1 ipsilateral to the active hand would be less readily attributable to direct motor execution and may instead reflect broader learning-related processes. Several prior studies have revealed hemispheric asymmetries in motor control, with the dominant left hemisphere in right-handed individuals showing broader involvement across effectors and a prominent role in organizing complex coordinated actions. During bimanual movements, for example, directed EEG coupling is biased from the dominant toward the nondominant sensorimotor cortex, suggesting greater influence of the left hemisphere over bilateral motor organization (Serrien et al., 2003). Consistent with this view, complex movements of the nondominant left hand recruit the ipsilateral left motor cortex more strongly than comparable right-hand movements recruit the right hemisphere (Koeneke et al., 2004), while learning a 90° rhythmic hand-foot coordination pattern engages a predominantly left-lateralized parietal-premotor network that is shared across left- and right-sided effector configurations (Swinnen et al., 2010). More recently, intracranial recordings showed that the left hemisphere encoded movement kinematics of both arms, whereas corresponding representations in the right hemisphere were largely restricted to the contralateral arm (Merrick et al., 2022). Neuropsychological evidence further supports asymmetric hemispheric contributions, in which left-hemisphere stroke disrupts trajectory formation and intersegmental coordination even in the ipsilesional arm (Haaland et al., 2009; Schaefer et al., 2007, 2009). Collectively, these findings support a model in which both hemispheres contribute to motor control but do so asymmetrically, with the dominant left hemisphere exerting broader influence over the organization and coordination of actions performed across different effector configurations (Serrien et al., 2006).

In the present study, corticospinal excitability in the dominant left M1 was expected to increase following both left- and right-hand unimanual training, reflecting its broader involvement across effectors. In contrast, modulation of the right M1 was expected to be more effector dependent, with greater changes following bimanual and left-hand training, when the right M1 directly contributes to movement execution. Consistent with our prediction, training-related corticospinal excitability showed a consistent positive change in the dominant left M1 across bimanual and both unimanual conditions, supporting its broader involvement across effector configurations and a possible contribution to representing or implementing the learned spatiotemporal coordination structure. Interestingly, right-M1 excitability also increased in all training conditions without clear dependence on the trained hand. Notably, the change in right-M1 excitability following right-hand training was not distinguishable from that observed during bimanual and left-hand training, when the right M1 was directly involved in movement control. Thus, the overall excitability profile was more bilateral than predicted and cannot be explained solely by the motor demands imposed on the contralateral M1.

The correlational findings further constrain the functional interpretation of these excitability changes. Greater increases in left-M1 excitability were associated with larger online gains in coordination accuracy only during right-hand training, whereas no corresponding relationship was observed during bimanual or left-hand training. Because the left M1 is directly involved in preparing and executing the active right-hand movement, this condition-specific association may reflect improved implementation of the motor commands required to coordinate the hand with the virtual partner rather than encoding of the 90° phase relationship itself. Moreover, greater increases in right-M1 excitability following right-hand training were associated with poorer offline improvements in accuracy and stability. The opposing behavioral associations of the two M1s raise the possibility that learning and subsequent memory stabilization depend partly on the relative functional balance between hemispheres rather than on excitability within either M1 alone (Duque et al., 2008; Paparella et al., 2023; Perez et al., 2007). The absence of comparable single-hemisphere associations during bimanual and left-hand training is also compatible with this interpretation, although it does not provide direct evidence for an interhemispheric balance mechanism. Overall, the limited and context-dependent relationship between corticospinal excitability and behavior provides weak evidence that M1 excitability directly reflects acquisition of the 90° spatiotemporal representation (Inchon Park et al., 2020). Instead, the opposing hemispheric associations raise the possibility that behavioral improvement depends more strongly on how activity is balanced and coordinated across the two M1s, motivating examination of interhemispheric interactions as a potential mechanism underlying coordination learning.

### Effector-dependent changes in interhemispheric inhibition

Building on the possibility that coordination learning depends on the relative organization of the two hemispheres, Experiment 3 examined practice-induced changes in IHI bidirectionally between the two M1s. IHI was assessed at a 40-ms interstimulus interval, corresponding to long-latency interhemispheric inhibition (IHI40). This later phase of IHI is mediated through transcallosal interactions that recruit inhibitory circuits in the opposite M1 and has been linked particularly to GABA_B-mediated intracortical inhibition (Irlbacher et al., 2007; Ni et al., 2009; Udupa et al., 2010). More broadly, modulation of IHI between the two M1s has been implicated in motor learning and the transfer of acquired motor skills across limbs (Paparella et al., 2023; Perez et al., 2007). Thus, bidirectional IHI provided a complementary measure to corticospinal excitability for determining whether learning the same spatiotemporal pattern reorganized the functional relationship between the two hemispheres.

In contrast to the broadly similar corticospinal excitability response across conditions, IHI revealed a task-specific pattern of interhemispheric reorganization. Bimanual training produced no reliable directional change in IHI, suggesting that simultaneous involvement of both M1s did not preferentially shift inhibitory influence toward either hemisphere. Unimanual training, however, produced asymmetric changes that depended on the practice hand. Following right-hand training, inhibition toward the right M1 increased, consistent with the established role of the M1 controlling the active hand in suppressing the opposite motor cortex during unilateral actions and limiting unwanted motor output (Duque et al., 2005; Fling & Seidler, 2011; Hübers et al., 2008). In contrast, left-hand training reduced inhibitory influence from the right M1 toward the dominant left M1. This pattern is counter to a strictly effector-dependent account, which would predict greater influence of the right M1 when it directly controls the active left hand. Instead, both unimanual conditions produced an interhemispheric organization that preserved the functional influence of the dominant left M1.

Several mechanisms may account for this apparent left-hemisphere bias. According to the hemispheric-specialization account, the dominant left hemisphere contributes broadly to organizing actions performed by either limb by exerting a stronger directional influence during bilateral coordination (Merrick et al., 2022; Serrien et al., 2003; Serrien et al., 2006; Swinnen et al., 2010). Under this account, reduced inhibition toward the left M1 following left-hand practice may reflect continued involvement of the dominant hemisphere in organizing the learned spatiotemporal structure, even when the right M1 directly controls the active hand. Relatedly, the proposal that a single hemisphere controls movement initiation for both hands (Stucchi & Viviani, 1993) provides a potential mechanism through which the dominant hemisphere could influence temporal organization across effector contexts. A complementary task-complexity account emphasizes that complex unilateral actions can recruit bilateral motor networks rather than only the M1 contralateral to the moving hand (Koeneke et al., 2004). Because the present task required continuous synchronization of the active hand with a virtual partner according to a 90° relationship, bilateral M1 involvement may also reflect the coordinative demands of implementing the learned pattern. Thus, the observed bias may arise from both dominant-hemisphere involvement in organizing skilled actions and bilateral motor recruitment during complex coordination.

Nevertheless, practice-induced changes in IHI were not reliably associated with either online or offline behavioral improvement. Although IHI was sensitive to the effector configuration through which the pattern was practiced, the magnitude of this reorganization did not predict how accurately or stably the pattern was learned. One possible explanation is that spatial and temporal coupling rely on partially distinct neural mechanisms (Heuer, 1993; Ivry et al., 2004). Interhemispheric communication through the corpus callosum appears particularly important for spatial interactions, as individuals with callosal disconnection exhibit less spatial interference than healthy controls while retaining temporal coupling (Franz et al., 1996). This persistent temporal coupling may instead be supported by a subcortical mechanism that accesses both effectors and gates the implementation of cortically generated motor commands (Ivry & Hazeltine, 1999; Ivry & Richardson, 2002). Because the defining requirement of the present 90° pattern was the temporal relationship between the limbs, IHI changes may reflect effector-dependent spatial interactions or motor implementation rather than learning of the temporal coordination structure itself. Consistent with this interpretation, experimentally modulating bilateral M1 excitability during learning of a 90° bimanual pattern affected movement amplitude more readily than relative-phase accuracy or its 6-hour retention (McCulloch et al., 2025), providing causal support for a preferential contribution of M1 to spatial implementation. Together, these findings suggest that M1 and its interhemispheric interactions contribute primarily to organizing and implementing the learned pattern, whereas its abstract temporal structure may depend on shared hemispheric or subcortical mechanisms.

### Motor network for encoding abstract spatiotemporal information

The limited relationship between M1 neurophysiological changes and coordination performance raises the possibility that the learned spatiotemporal relationship is represented within a broader motor network rather than within M1 alone. Neuroimaging studies of coordinated actions provide substantial support for this distributed account. Debaere et al. (2004) independently manipulated movement frequency and spatiotemporal complexity across unimanual and bimanual coordination patterns. Activity in the supplementary motor area and superior parietal cortex increased primarily with spatiotemporal complexity, whereas M1 activity was more closely related to movement frequency (Debaere et al., 2004). The sensitivity of dorsal premotor cortex and cerebellum to both spatiotemporal complexity and movement frequency suggests that these regions may bridge the representation of the coordination structure with its motor implementation.

Learning a novel coordination pattern similarly produces distributed changes across cortical and subcortical regions. During acquisition of a 90°bimanual pattern, learning-related modulation has been observed in premotor and parietal cortices, M1, cingulate motor areas, basal ganglia, thalamus, and cerebellum, consistent with a progressive redistribution of neural resources as performance becomes more skilled (Debaere et al., 2004). With extensive practice, Puttemans et al. (2005) further showed that increases in M1 activity observed during learning returned toward pretraining levels once performance became automated, whereas elevated activity in the putamen and anterior cerebellum persisted (Puttemans et al., 2005). This temporal dissociation suggests that M1 engagement may be particularly prominent during acquisition and implementation of the developing skill, whereas cerebellar and basal-ganglia circuits may contribute more persistently to the stabilization, long-term retention, and automatic expression of acquired motor skills (Doyon et al., 2009; Doyon & Benali, 2005; Hadjiosif et al., 2024; Puttemans et al., 2005).

Importantly, the present findings do not exclude the contribution of M1 to the acquisition of the 90° relative phase pattern. M1 activity changes during learning of novel coordination patterns, and its contribution may vary across different stages of skill acquisition (Debaere et al., 2004; Puttemans et al., 2005). Moreover, the present study assessed neurophysiological changes associated with learning but did not directly perturb M1 function. Accordingly, the weak and context-dependent associations between M1 excitability and behavioral improvement cannot establish that M1 is unnecessary for acquiring the 90°relationship. Recent bilateral M1 stimulation work similarly showed that altering the relative excitability of the two M1s had little effect on the accuracy of the 90° relative-phase during training or after a 6-hour retention, despite influencing movement amplitude, further suggesting that M1 modulation may affect motor implementation more readily than the spatiotemporal relationship itself (McCulloch et al., 2025). Future studies using temporally targeted inhibitory stimulation of M1 during practice will therefore be necessary to determine whether M1 is causally involved in acquiring novel relative-phase patterns.

## Conclusion

In conclusion, learning a novel 90° relative-phase pattern showed a common behavioral signature across bimanual and unimanual coordination, consistent with a shared, partially effector-independent representation of the newly acquired attractor in the context of coordination dynamics theory of motor control and learning (Buchanan et al., 2007). At the neurophysiological level, however, corticospinal excitability and interhemispheric inhibition exhibited distinct patterns that were not consistently associated with behavioral improvement, indicating that acquisition of the coordination pattern is not straightforwardly reflected in changes measured within and between the two M1s. Together, these findings highlight a dissociation between behavioral learning and M1 neurophysiological plasticity and point to the need to determine how M1 interacts with broader neural systems to support the acquisition and expression of novel spatiotemporal coordination patterns.

## Author Contributions

Y.W. contributed to the conceptualization, methodology, investigation, data analysis, and writing of the original draft. D.M.K. and J.J.B. contributed to the conceptualization, supervision, project administration, and critical revision of the manuscript. M.M.W. and Y.W. contributed to data collection, data curation, and formal analysis. D.L.W. and Y.L. contributed to the methodology, supervision, interpretation of results, and critical revision of the manuscript. All authors reviewed and approved the final version.

## Declaration of Generative AI and AI-Assisted Technologies

During the preparation of this work, the author(s) used Chat GPT to proofread the writing. After using this tool/service, the author(s) reviewed and edited the content as needed and take(s) full responsibility for the content of the publication.

