## Supplementary Tables for "Characterizing bilateral M1 excitability changes after learning a novel spatiotemporal coordination pattern"

Supplementary Material

Table 1. Correlations between ΔMEP (post − pre) and training-related improvement across groups

| Group | Tested M1 | ΔBehavior | n | Spearman's ρ | p | Pearson's r | p |
|---|---|---|---|---|---|---|---|
| BM | R | BW30 | 13 | 0.115 | .710 | 0.006 | .984 |
| BM | L | BW30 | 13 | -0.533 | .064# | -0.471 | .104 |
| BM | R | Variability | 13 | -0.165 | .591 | -0.066 | .830 |
| BM | L | Variability | 13 | 0.500 | .085# | 0.564 | .045* |
| UR | R | BW30 | 13 | 0.423 | .152 | 0.441 | .131 |
| UR | L | BW30 | 13 | 0.610 | .030* | 0.389 | .188 |
| UR | R | Variability | 13 | -0.374 | .209 | -0.304 | .313 |
| UR | L | Variability | 13 | -0.330 | .272 | -0.248 | .414 |
| UL | R | BW30 | 12 | -0.490 | .110 | -0.289 | .363 |
| UL | L | BW30 | 12 | 0.217 | .499 | 0.167 | .604 |
| UL | R | Variability | 12 | 0.441 | .154 | 0.677 | .016* |
| UL | L | Variability | 12 | 0.098 | .766 | 0.314 | .320 |

Note. Spearman correlation was treated as the primary analysis because of the modest sample size and its greater robustness to outliers, non-normality, and nonlinearity; Pearson correlations are provided as a supplementary analysis. ΔMEP was calculated as post − pre. * $p < .05$. # $p < .10$. BM = bimanual group; UR = unimanual right group; UL = unimanual left group.

Table 2. Correlations between ΔMEP (post − pre) and consolidation-related improvement across groups

| Group | Tested M1 | ΔBehavior | n | Spearman's ρ | p | Pearson's r | p |
|---|---|---|---|---|---|---|---|
| BM | R | BW30_6h | 13 | 0.033 | .921 | 0.057 | .855 |
| BM | L | BW30_6h | 13 | 0.308 | .306 | 0.444 | .128 |
| BM | R | Variability_6h | 13 | 0.016 | .964 | 0.179 | .558 |
| BM | L | Variability_6h | 13 | -0.027 | .935 | -0.043 | .888 |
| UR | R | BW30_6h | 13 | -0.698 | .010* | -0.769 | .002* |
| UR | L | BW30_6h | 13 | -0.549 | .055# | -0.327 | .276 |
| UR | R | Variability_6h | 13 | 0.896 | <.001* | 0.868 | <.001* |
| UR | L | Variability_6h | 13 | 0.462 | .115 | 0.404 | .171 |
| UL | R | BW30_6h | 12 | 0.196 | .543 | 0.174 | .589 |
| UL | L | BW30_6h | 12 | -0.21 | .513 | -0.373 | .233 |
| UL | R | Variability_6h | 12 | -0.42 | .177 | -0.276 | .385 |
| UL | L | Variability_6h | 12 | -0.126 | .700 | -0.032 | .921 |

Note. Spearman correlation was treated as the primary analysis because of the modest sample size and its greater robustness to outliers, non-normality, and nonlinearity; Pearson correlations are provided as supplementary analyses. ΔMEP was calculated as post − pre. ΔBehavior was calculated as the change from post-test to 6 h retention. * $p < .05$. # $p < .10$. BM = bimanual group; UR = unimanual right group; UL = unimanual left group.

Table 3. Correlations between ΔIHI (post − pre) and training-related improvement across groups

| Group | Tested M1 | ΔBehavior | n | Spearman's ρ | p | Pearson's r | p |
|---|---|---|---|---|---|---|---|
| BM | R | BW30 | 12 | 0.238 | .457 | 0.316 | .317 |
| BM | L | BW30 | 12 | 0.049 | .886 | -0.034 | .917 |
| BM | R | Variability | 12 | -0.49 | .110 | -0.53 | .076# |
| BM | L | Variability | 12 | -0.308 | .331 | -0.075 | .818 |
| UR | R | BW30 | 12 | -0.252 | .430 | -0.07 | .830 |
| UR | L | BW30 | 12 | -0.203 | .528 | -0.348 | .268 |
| UR | R | Variability | 12 | -0.252 | .430 | -0.026 | .935 |
| UR | L | Variability | 12 | 0.126 | .700 | 0.26 | .415 |
| UL | R | BW30 | 12 | -0.147 | .651 | -0.016 | .961 |
| UL | L | BW30 | 12 | 0.014 | .974 | 0.267 | .402 |
| UL | R | Variability | 12 | -0.098 | .766 | -0.224 | .484 |
| UL | L | Variability | 12 | -0.371 | .237 | -0.382 | .221 |

Note. Spearman correlation was treated as the primary analysis because of the modest sample size and its greater robustness to outliers, non-normality, and nonlinearity; Pearson correlations are provided as supplementary analyses. ΔIHI was calculated as post − pre. ΔBehavior was calculated as the change from pre-test to post-training. * $p < .05$. # $p < .10$. BM = bimanual group; UR = unimanual right group; UL = unimanual left group.

Table 4. Correlations between ΔIHI (post − pre) and consolidation-related improvement across groups

| Group | Tested M1 | ΔBehavior | n | Spearman's ρ | p | Pearson's r | p |
|---|---|---|---|---|---|---|---|
| BM | R | BW30 | 12 | -0.077 | .817 | -0.047 | .883 |
| BM | L | BW30 | 12 | -0.427 | .169 | -0.389 | .211 |
| BM | R | Variability | 12 | 0.245 | .444 | 0.152 | .637 |
| BM | L | Variability | 12 | 0.301 | .343 | 0.279 | .380 |
| UR | R | BW30 | 12 | 0.070 | .834 | 0.167 | .605 |
| UR | L | BW30 | 12 | 0.000 | 1.000 | 0.043 | .893 |
| UR | R | Variability | 12 | -0.210 | .513 | -0.334 | .288 |
| UR | L | Variability | 12 | -0.168 | .604 | 0.026 | .936 |
| UL | R | BW30 | 12 | 0.259 | .417 | 0.461 | .132 |
| UL | L | BW30 | 12 | -0.252 | .430 | -0.174 | .589 |
| UL | R | Variability | 12 | -0.357 | .256 | -0.493 | .103 |
| UL | L | Variability | 12 | 0.378 | .227 | 0.203 | .526 |

**Note.** Spearman's rank correlation was treated as the primary analysis because of the modest sample size and its greater robustness to outliers, non-normality, and nonlinearity; Pearson correlations are reported as supplementary analyses. ΔIHI was calculated as post-training minus pre-training IHI. Behavioral measures were assessed at the 6-hour retention test and included performance within ±30° of the target coordination pattern (BW30) and coordination variability. $p < .05$. #$p < .10$. BM = bimanual group; UR = unimanual right group; UL = unimanual left group.
